# Eye movements in real-life search are guided by task-irrelevant working-memory content

**DOI:** 10.1101/2020.05.18.101410

**Authors:** Cherie Zhou, Monicque M. Lorist, Sebastiaan Mathôt

**Author notes:** Correspondence concerning this article should be addressed to Cherie Zhou, Department of Experimental Psychology, University of Groningen, Grote Kruisstraat 2/1, 9712 TS, Groningen.

## Abstract

Attention is automatically guided towards stimuli that match the contents of working memory. This has been studied extensively using simplified computer tasks, but it has never been investigated whether (yet often assumed that) memory-driven guidance also affects real-life search. Here we tested this open question in a naturalistic environment that closely resembles real life. In two experiments, participants wore a mobile eye-tracker, and memorized a color, prior to a search task in which they looked for a target word among book covers on a bookshelf. The memory color was irrelevant to the search task. Nevertheless, we found that participants’ gaze was strongly guided towards book covers that matched the memory color. Crucially, this memory-driven guidance was evident from the very start of the search period. These findings support that attention is guided towards working-memory content in real-world search, and that this is fast and therefore likely reflecting an automatic process.

**Significance statement:** A core concept in the field of visual working memory (VWM) is that visual attention is automatically guided towards things that resemble the content of VWM. For example, if you hold the color red in VWM, your attention and gaze would automatically be drawn towards red things in the environment. So far, studies on such memory-driven guidance have only been done with well-controlled computer tasks that used simplified search displays. Here we address the crucial and open question of whether attention is guided by the content of VWM in a naturalistic environment that closely resembles real life. To do so, we conducted two experiments with mobile eye tracking. Crucially, we found strong memory-driven guidance from the very early phase of the search, reflecting that this is a fast, and therefore likely automatic, process that also driven visual search in real life.

---

Searching is a complex task. When you are looking for an object in your environment, you generally keep a prototype of the object in working memory; this prototype can be defined by different stimulus attributes (e.g., shape, color, size, name; Wolfe, 1994). The visual system efficiently directs our attention towards objects that resemble the features stored in working memory, a phenomenon referred to as memory-driven guidance of attention (Bundesen, 1990). Theories of visual working memory (VWM) and attention explain this by positing that the neural representations of visual features that are maintained in VWM are pre-activated (Chelazzi et al., 1993; Desimone, 1998); therefore, when the incoming visual information matches the features that are maintained in VWM, the representation of these matching features is boosted by virtue of riding on top of this pre-activation, thus resulting in attentional guidance towards these features.

Memory-driven guidance has been extensively investigated in lab studies. For example, in a combined memory-search paradigm (Olivers, Meijer, & Theeuwes, 2006), participants were first asked to keep a color in memory. During the retention interval, they performed a visual search task in which the search display consisted of: a target; a singleton distractor that matched or did not match the memory color; and several gray distractors. Finally, in a memory test, participants indicated the memorized color among three colored disks. Importantly, Olivers and colleagues found that the interference of the singleton distractor, as measured through increased response times (RTs) to the target, was especially high when its color matched the memory color maintained in working memory.

Jung and colleagues (2018) used real-world images in a combined memory-search task. Their task was similar to the one used by Olivers et al. (2006), but more closely resembled real-world search. (Although it was still a computer task.) Participants were first asked to keep an object in memory (e.g., a coffee cup). Next, they searched for a target in indoor-scenes images (e.g., a photo of a room). In line with previous findings, Jung and colleagues found that search was slowed when the memorized object appeared in the search scene as a distractor, presumably because attention was automatically (mis)guided towards the memorized object.

So far, evidence for memory-driven guidance has only come from well-controlled computer tasks (e.g., Bahle, Beck, & Hollingworth, 2018; Soto, Heinke, Humphreys, & Blanco, 2005). Yet the assumption is that memory-driven guidance is a fundamental mechanism that allows us to search for things in real life. Can we still find robust memory-driven guidance when stimuli do not appear only at a limited set of locations on a computer screen, but appear anywhere in a real-life environment, while participants are free to move and look wherever they want? This question is crucial because the few previous studies of real-life visual search (on topics other than memory-driven guidance) have shown that some cognitive effects that are observed in the lab do not transfer to real life (e.g., Brennan, Watson, Kingstone, & Enns, 2009).

In the present study, we created a naturalistic search setting that resembles searches in daily life far more closely than has so far been done in previous studies. Participants wore a mobile eye-tracker, and looked for a target word among book covers while maintaining a color in working memory. As a search array, we used real bookshelves with book covers in four different color categories: Blue, Yellow, Red, and Green. We predicted that, if attention is guided towards the color that is maintained in VWM, participants would look more often at books that match the memory color, even though color is irrelevant to the search task. Moreover, if the color of the book cover that contains the target word matches the memory color, the search RT should be faster. Consistent with our prediction, the eye-tracking data for two experiments demonstrated a very strong guidance effect in this real-world search task. There was also a tendency for a memory-driven guidance effect in RTs.

## Method

### Preregistration

A detailed pre-registration of Experiment 2 is available at https://osf.io/nxbzh. All deviations from the preregistration will be mentioned below.

### Participants

Fifteen and twenty-six first-year psychology students from the University of Groningen participated in Experiment 1 and 2, respectively, in exchange for course credits. (This deviates from the preregistered 45 participants for Experiment 2. We decided to stop data collection when we found that the memory-driven-guidance effect was unexpectedly strong, such that a smaller number of participants already provided sufficient statistical power.) All participants had normal or corrected-to-normal acuity and color vision. The studies were approved by the local ethics review board of the University of Groningen (PSY-1819-S-0224; PSY-1920-S-0115). Participants provided written informed consent before the start of the experiment.

### Setup

Participants were equipped with a wearable eye-tracker (Pupil labs, Berlin, Germany). Eye movements of the right eye were recorded with an eye camera on the eye-tracker. A video from the participant’s view was recorded with a scene camera, at a sampling rate of 14 Hz (Exp 1) and 30 Hz (Exp 2). (Although the eye camera has a higher sampling rate, the sampling rate of the scene camera determined the temporal resolution of our analysis.) In Experiment 1, we calibrated the eye-tracker using “Manual Marker Calibration” provided in the Pupil Capture software (version 1.11.4; Kassner et al., 2014). Participants stood in front of the experimenter (CZ) at a distance of approximately 1.5m, and gazed at a printed marker that was presented at nine locations by CZ. In Experiment 2, we calibrated the eye-tracker using “Screen Marker Calibration” provided in the Pupil Capture software (version 1.18; Kassner et al., 2014). Participants sat behind a computer screen at a viewing distance of approximately 45 cm, and gazed at a marker that was presented at five locations on the screen. Four printed square markers surrounded each book on the shelf (*Figure 1*); these markers were recognized automatically by the Pupil Player software and allowed us to define a region of interest (ROIs) for each book in real-world coordinates.

**Figure 1.**
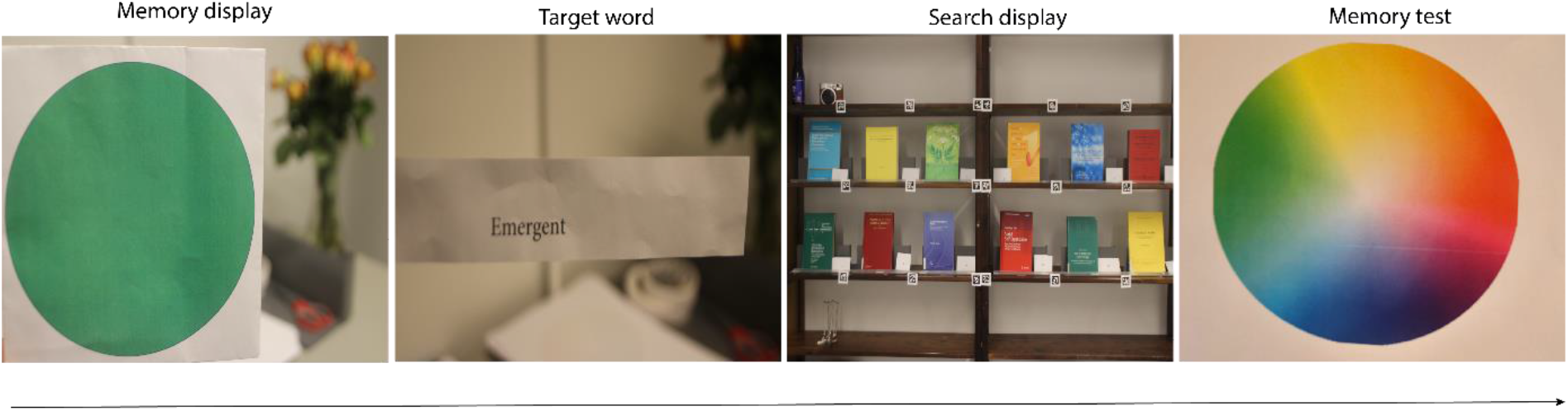
Sequence of events in Experiment 2. Participants first saw a color that they kept in visual working memory. Next, they saw a target word, and searched for a book cover that contained this word. Finally, they indicated the memorized color on a color circle.

### Stimuli, design and procedure

In Experiment 1, participants were instructed to memorize the exact shade of a color (*Figure 1*), for later recall in a memory test. Each memory color was selected from a color category: blue, yellow, red, or green (Memory Color). Next, participants searched for a target word among 16 book covers on a bookshelf, during which gaze data was collected. Four book covers were selected from each color category: blue, yellow, red, or green (Book Color). Participants were told to move around freely while they searched. The book covers had different amounts of text and different font sizes, and were of different languages (English, Dutch, German, and Polish). One target word was selected from each book cover. Participants said the book number out loud as soon as they found the target word and looked away from the shelf. Their response times (RTs) were recorded manually using a stopwatch by CZ. Each trial ended with a memory test, in which participants indicated the exact shade of the memory color by drawing a line or a cross on a printed color circle. Feedback on the memory test followed, allowing participants to compare the color that they had indicated with the color that they had seen. Each participant completed eight trials. On some trials, the book cover that contained the target word belonged to the same color category as the memory color (Target-match), and on other trials it belonged to a different color category (Target-non match). The Target Book Color was either in the same color category as (Target-match), or in a different color category from (Target-non-match), the Memory color.

In Experiment 2, the procedure was the same as in Experiment 1 except for the following. In the search task, participants searched for a target word among 12 book covers in a bookshelf, at a distance of at least 71 cm in order for the eye-tracker to detect the square markers. All book covers had a similar amount of text and a similar font size, and all books were English. The memory colors and target words were selected pseudorandomly such that each color category occurred twice. After each search, participants were told of their RT as feedback to encourage fast responses.

### Data processing

The ROI of each book was defined using the “Offline Surface Tracker” (version 1.11.4; Kassner et al., 2014) in Experiment 1 and the “Surface Tracker” plugin in Experiment 2 in the Pupil Player software (version 1.18; Kassner et al., 2014) with four square markers. Gaze position was detected by the infrared camera using the “dark pupil” method, and gaze samples on each surface were exported. The timeframes for the start point and end point of each trial were coded based on a visual inspection of the video by CZ. When gaze position seemed to be biased (e.g., systematically just below a book cover), this was manually corrected in a separate document. Gaze Proportion on each book color (i.e. Blue, Yellow, Red and Yellow) relative to the total number of samples of each trial was calculated.

If more than 20% gaze points or video frames (due to technical malfunction) were lost, or if 20% of gaze points deviated so much from a surface that they could not be manually corrected, the trial was excluded from the eye-tracking analysis. (This is slightly different from the preregistration, in which we did not specify the proportion of lost data, and did not include lost video frames as a criterion for exclusion.) When participants forgot or failed to find the target word, or when the search was incorrect, the trial was excluded from both eye-tracking and behavioral analysis. (This criterion was not preregistered. We added it to exclude inaccurate search trials.) In Experiment 1, if the participants reported that their search was biased because the target matched their native language, the trial was also excluded from both analyses. Moreover, trials with response times shorter than 1s or longer than 120s were excluded from the behavioral analysis. In total, 96 trials (of 119) and 157 trials (of 207) remained for the eye-tracking analysis, and 111 trials (of 119) and 198 trials (of 207) remained for the behavioral analysis in Experiment 1 and 2, respectively.

### Statistical analyses

#### Eye-tracking analysis

We conducted linear mixed effects models (LMER) using the R package lmerTest using *Gaze Proportion* as dependent measure, *Memory Color* and *Book Color* (Blue, Yellow, Red, Green) as fixed effects, with a random intercept by participant, and a random intercept by trial which was nested in participant. We built two models: in the first model (lm1), the variance was accounted for by the main effect of Memory Color and Book Color; in the second model (lm2), we added the interaction between the two effects. Next, we determined which model better accounted for the data by comparing the χ^2^ and the associated p-value from the log-likelihood of the two models (Baayen et al., 2008).

#### Behavioral analysis

We conducted a LMER using *Reaction Time* as dependent measure, *Target Match* (Target-match, Target-non-match) as fixed effect, with a random by-participant intercept and a random by-participant slope for the effect of Target.

## Results

As shown in Figure 2, when participants maintained a color in working memory, they looked more at the book covers that matched the memory color during the search task, as compared to book covers that did not match the memory color, in both Experiment 1 (Memory Color × Book Color: χ^2^ (9, n= 15) = 39.03, p= 1.14 ×10^-5^; Figure 2a) and Experiment 2 (χ^2^ (9, n= 23) = 71.69, p= 7.08 ×10^-12^; Figure 2b). Importantly, this effect was already present from the very start of the search trial and remained constant over time (Figure 3), consistent with the notion that memory-driven guidance of attention is a fast and automatic process. We did not find a reliable effect of Target Match on RTs, although it was in the predicted direction for both experiments (Exp 1: M(Target-match) = 33.69s, M(Target-non-match) = 40.31s; *t* = −1.01, *p* = 0.33; Exp 2: M(Target-match) = 25.58s, M(Target-non-match) = 32.86s; *t* = −1.89; *p*= 0.07).

**Figure 2.**
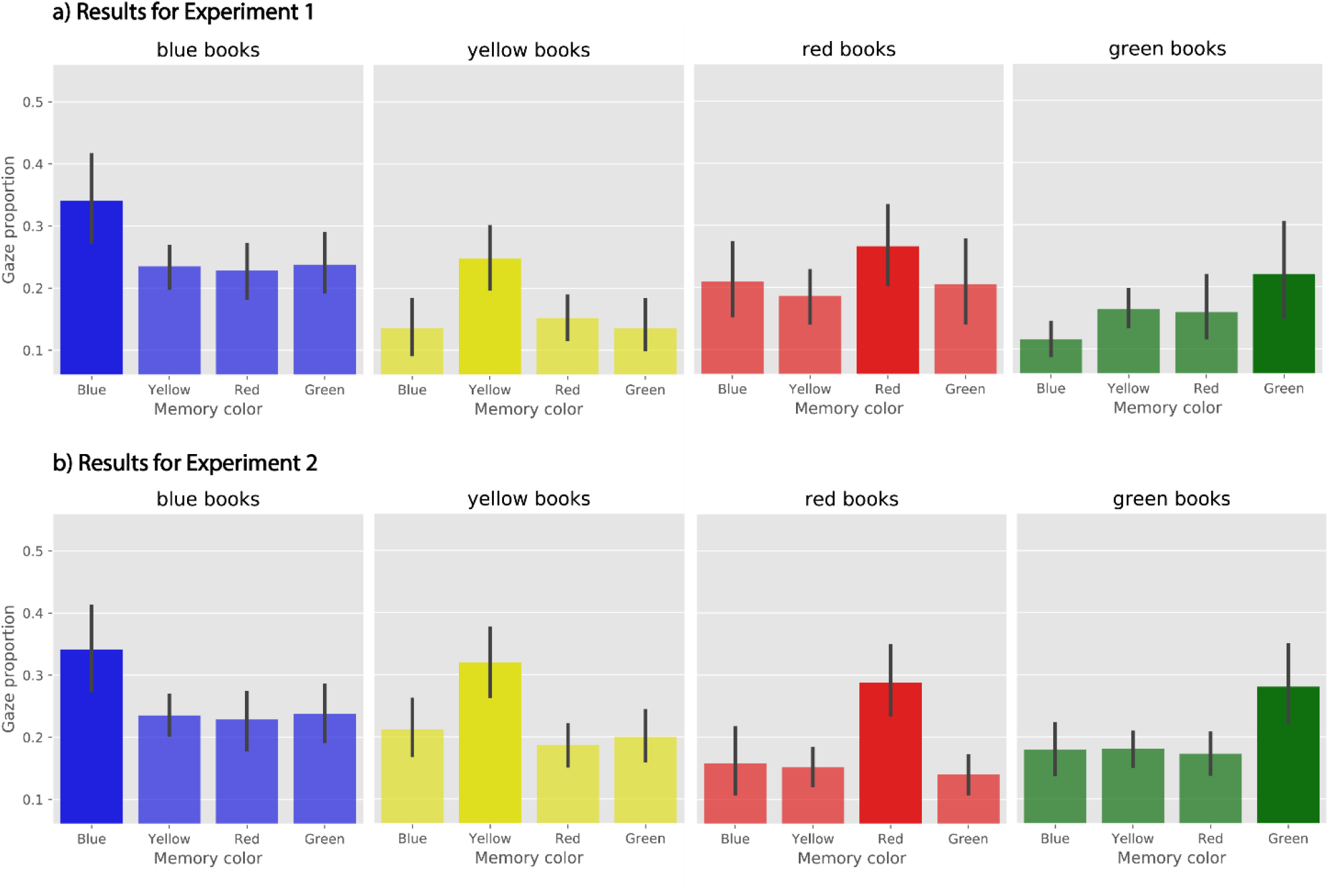
Gaze proportion on book covers in four color categories (Blue, Yellow, Red and Green) as a function of memory color (Blue, Yellow, Red and Green) in Experiment 1 (a) and Experiment 2 (b).

**Figure 3.**
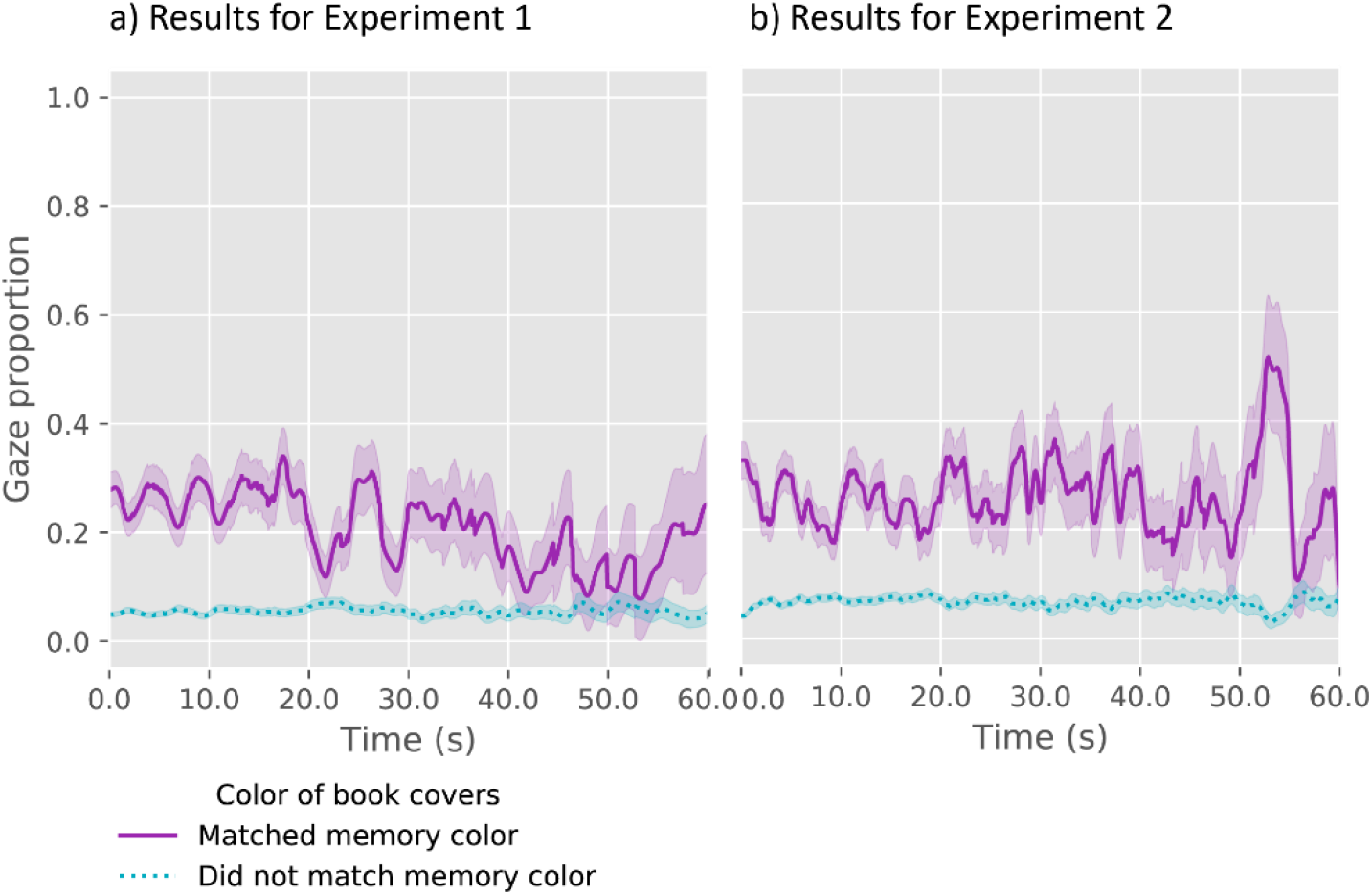
Gaze proportion on book covers that matched (i.e. matching book covers; purple solid line) or did not match (i.e. non-matching book covers; teal dotted line) the memory color in a search trial over time in Experiment 1 (a) and Experiment 2 (b). The results for the non-matching book covers reflects the average gaze proportion over the three non-matching covers.

## Discussion

The present study shows that visual working memory guides attention in real-world visual search. In two experiments, participants wore an eye-tracker while performing a combined memory-search task in a naturalistic setting. First, participants memorized a color shade that they had to report later. Next, they searched for a target word among book covers that matched or did not match the color in working memory. We found that participants’ gaze was guided towards the book covers that matched the memory color during visual search, as compared to the covers that did not match the memory color. Moreover, this effect was present from the very beginning of the search, remained constant over time, and occurred despite the fact that the memory color was irrelevant for the search task.

Unlike most lab-based studies that generally use simplified search stimuli, visual search in real-life environments is affected by distractor objects that vary strongly in their visual characteristics and locations. In Experiment 1, we used book covers that had different features (e.g., font sizes, amounts of text, languages), which created a natural setting that closely resembled a real-world situation. In contrast, in Experiment 2, we created a more controlled environment with more unified features (e.g., similar font size and amount of text, all texts in English). Consistent with previous findings (Chen & Zelinsky, 2006; Seidl-Rathkopf, Turk-Browne, & Kastner, 2015), the results of Experiment 1 and 2 suggest that in real-life search, VWM content provide a top-down template that guides attention, regardless of whether it is in a highly variable (Experiment 1) or more controlled (Experiment 2) environment.

Although we did not find a reliable memory-driven guidance effect on reaction times (RTs), both experiments showed a tendency towards faster RTs when the book color of the target word matched the memory color, as we predicted. Our real-life search task provided only a small number of trials (eight) for each participant, as compared to computer tasks in which each participant performs hundreds of trials. This likely provided insufficient statistical power for analyzing the behavioral results. In contrast, the lengthy searches provided us with a lot of eye-movement data, and therefore excellent statistical power for analyzing the eye-movement results.

To conclude, our results provide an important extension of previous findings (Bahle et al., 2018; Jung et al., 2018; Soto et al., 2005) by showing that VWM content guides attention in real-world search, even when it is irrelevant to the search task. Crucially, this effect appeared already at the very early stage of visual search, showing that attention is automatically driven toward working memory contents (Seidl-Rathkopf et al., 2015; Soto et al., 2005, 2008)—not only on a computer screen, but also when you’re looking for that lost set of keys in your living room.

